# Genetic variations within human gained enhancer elements affect human brain sulcal morphology

**DOI:** 10.1101/2021.09.10.459622

**Authors:** Herve Lemaitre, Yann Le Guen, Amanda K. Tilot, Jason L. Stein, Cathy Philippe, Jean-François Mangin, Simon E. Fisher, Vincent Frouin

## Abstract

The expansion of the cerebral cortex is one of the most distinctive changes in the evolution of the human brain. Cortical expansion and related increases in cortical folding may have contributed to emergence of our capacities for high-order cognitive abilities. Molecular analysis of humans, archaic hominins, and non-human primates has allowed identification of chromosomal regions showing evolutionary changes at different points of our phylogenetic history. In this study, we assessed the contributions of genomic annotations spanning 30 million years to human sulcal morphology measured via MRI in more than 18,000 participants from the UK Biobank. We found that variation within brain-expressed human gained enhancers, regulatory genetic elements that emerged since our last common ancestor with Old World monkeys, explained more trait heritability than expected for the left and right calloso-marginal posterior fissures and the right central sulcus. Intriguingly, these are sulci that have been previously linked to the evolution of locomotion in primates and later on bipedalism in our hominin ancestors.

## Introduction

“Nothing in biology makes sense except in the light of evolution” is one of the most cited declarations in biology (Dobzhansky, 1973) and the brain does not seem to escape this rule. Indeed, one way to better understand the structures and functions of the human brain in its current form is to understand their evolutionary history. Modern human brains were shaped by the cumulative effects of mutation, genetic recombination, and other sources of genetic variation over millions of years of evolution. While genetic variations that shaped the brain across hominin evolution may have facilitated adaptation of humans to their environment, it is thought that some may have also increased the risk for neurodevelopmental and neurodegenerative disorders (Bruner & Jacobs, 2013; Bufill et al., 2011).

Some of the most salient features of human brain evolution have been the expansion of the endocranium and a shift towards a distinctive globular shape (Gunz et al., 2019; Holloway et al., 2009). Increased skull size is thought to be driven in part by an enlargement of the cerebral cortex (Rakic, 2009). The cortex has not expanded uniformly; instead, specific cortical areas have expanded more than others, which has also influenced cortical folding of nearby regions.

Cortical folding consists of the formations of gyri and their counterparts called sulci. This feature of neuroanatomy has often been considered in regard to evolution and ontogeny, as it varies widely among mammals and originates early during brain development (Zilles et al., 2013). Since cortical folding starts very early in life and occurs in a precise spatio-temporal order (Kersbergen et al., 2016), it has been hypothesized to be under strong genetic control (Le Guen et al., 2018). The central sulcus, one of the most prominent cortical folds, shows changes in surface area, shape and folding patterns during Old World monkey evolution (around 30 million years) until the emergence of anatomically modern humans (Hopkins et al., 2014). Changes in cortical folding patterns may have contributed to the emergence of our capacities for high-order cognitive abilities such as creativity and language. However, specific insights into the evolutionary history of the human brain are lacking due to its fast decomposition after death. Indeed, the direct study of this question has been impossible due to the absence of fossilized brain tissue from our hominin ancestors.

Comparative brain studies with extant primates, related to humans through common ancestors, have provided an indirect way to reconstruct some of the major changes that the brain underwent on the lineage that led to Homo sapiens (Heuer et al., 2019). Recent progress in the field of paleogenomics has given opportunities to study human brain evolution from a novel perspective (Pääbo, 2014). Molecular analyses of multiple human and non-human primate populations, along with DNA from fossils of archaic hominins, has made it possible to use comparative genomic and population genetic annotations to identify evolutionarily relevant loci in the human genome across diverse time scales, including: Neanderthal introgression – ≈ 50 kya - (Vernot et al., 2016), selective sweeps – ≈ 250 kya - (Peyrégne et al., 2017), human accelerated regions - ≈ 7 Mya - (Capra et al., 2013; Vermunt et al., 2016) and human gained enhancers – ≈ 30 Mya - (Reilly et al., 2015; Vermunt et al., 2016). Moreover, using large-scale neuroimaging genetic approaches, we can now study the relationship between evolutionarily annotated genetic variants and interindividual variability in brain phenotypes in living humans. These approaches have been used with success to study surface area and thickness of different cortical regions (Tilot et al., 2020) as well as measures of functional connectivity (Wei et al., 2019). Wei et al. found an association between human accelerated regions and functional activity within the default mode network in data from 6,899 participants from the UK Biobank (Sudlow et al., 2015). Tilot et al. reported association between human gained enhancers and surface area in **several** regions, including the inferior frontal gyrus, in a sample of more than 30,000 individuals from the Enigma consortium (Grasby et al., 2020) including data from the UK Biobank.

In the present study, we aimed to expand this understanding of human brain evolution by assessing the relationship of genomic annotations spanning 30 million years with interindividual differences in human sulcal morphology in more than 18,000 participants from the UK Biobank. We hypothesized that genetic variation affecting interindividual differences in sulcal features may be enriched in genomic regions under selective pressure or those which have gained functionality over human evolution, which would demonstrate that variations within these evolutionary elements are important for regulating the areal organization of the human cortex.

## Methods

The analyses described here were conducted under UK Biobank data application number 25251. The UK Biobank is a health research resource that aims to improve the prevention, diagnosis, and treatment of a wide range of illnesses. Between the years 2006 and 2010, about 500,000 people aged between 45 and 73 years old were recruited in the general population across Great Britain. In this work, we used the T1-weighted MRI data available as of March 2018 from 20,060 subjects. The imaging quality control was performed by UK Biobank (Alfaro-Almagro et al., 2018).

The cortical sulci were extracted from T1-weighted images using BrainVISA (http://brainvisa.info). Morphologist 2015, an image processing pipeline included in BrainVISA, was used to quantify sulcal parameters. Briefly, the Morphologist 2015 segmentation pipeline extracts left and right hemisphere masks, performs gray and white matter classification, and computes a negative cast of each cortical fold (J. F. Mangin et al., 2004). Then, a Bayesian pattern recognition approach relying on a probabilistic atlas is used to label the folds using a nomenclature of 125 sulci (Perrot et al., 2011). The sulcus recognition process combines localization and shape information. The atlas is described in detail and freely accessible here: http://brainvisa.info/web/morphologist.html and can be visualized online here: http://brainvisa.info/web/webgl_demo/webgl.html. For each sulcus, sulcal opening and depth features were computed. Sulcal opening was defined as the average distance between both banks of the pial surface. Sulcal depth was defined as the average geodesic distance from the convex hull of the brain to the bottom line of the sulcus medial surface (J.-F. Mangin et al., 2010). In prior investigations of sulcal opening measures and single-nucleotide polymorphism (SNP) DNA data from the UK Biobank, twenty two main brain sulci were found to show significant SNP-based heritability (Le Guen et al., 2019). We therefore selected those twenty-two sulci for analysis in the current study (Figure 1).

**Figure 1.**
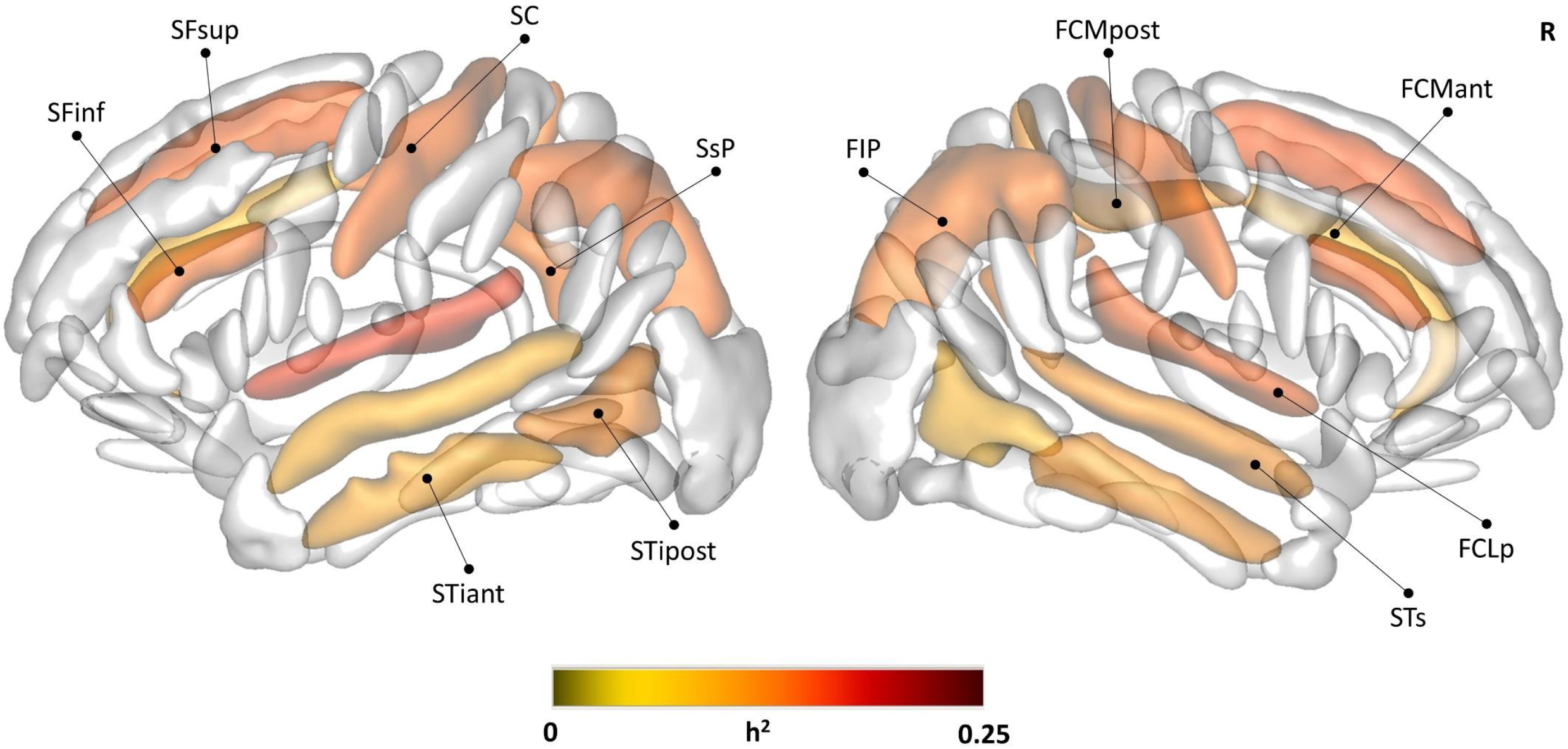
SNP-based heritability estimates of sulcal opening for the twenty-two sulci. Bonferroni corrected *P*-value< 0.5. FCLp: posterior lateral fissure; FCMant: calloso-marginal anterior fissure; FCMpost: calloso-marginal posterior fissure; FIP: intraparietal sulcus; SC: central sulcus; SFinf: inferior frontal sulcus; SFsup: superior frontal sulcus; STiant: anterior inferior temporal sulcus; STipost: posterior inferior temporal sulcus; STs: superior temporal sulcus; SsP: sub-parietal sulcus. The sulci are displayed using the statistical probability anatomy map (SPAM) representation, which represents the average sulci shape and position on the reference base of the Brainvisa sulci extraction pipeline (Perrot et al. 2011), h^2^: SNP-based heritability, R: Right.

Genotyping in the UK Biobank was performed using the UK BiLEVE Axiom array by Affymetrix on a subset of 49,950 participants (807,411 markers) and the UK Biobank Axiom array on 438,427 participants (825,927 markers). Arrays are similar and share 95% of common SNP probes. The imputed genotypes were obtained from the UK-BioBank repository and consisted of 93,095,623 autosomal SNPs using the Haplotype Reference Consortium and UK10K + 1000 Genomes reference panels (Bycroft et al., 2017). We limited our analysis to people identified by UK Biobank as belonging to the main white British ancestry subset. Additionally, we relied on the Quality Control carried out by the UK Biobank consortium on the genotyping data which excluded participants with high missingness, high heterozygosity, first degree related individuals or sex mismatches (Bycroft et al., 2017). In total 18,101 participants passed the image processing steps and the genetic criteria filtering (8547 females, 62.75 ± 7.45 years).

Genome-wide association studies (GWAS) were performed assessing association between the filtered imputed genotypes and the sulcal opening and depth of the eleven left and eleven right selected sulci, using PLINK 1.9 (Purcell et al., 2007) with the following thresholds: missing genotypes < 10%, Hardy-Weinberg Equilibrium p < 10^−6^ and minor allele frequency < 1.0% for a total of 8,376,022 autosomal SNPs tested. Sex, age, the genotyping array type and the first 10 genetic ancestry principal components were added as covariates.

The contributions of each SNP set (defined based on evolutionary annotations) to the total SNP heritability of each sulcal trait were determined using partitioned heritability analyses from the previously obtained GWAS summary statistics, following methods described in Tilot et al. 2020 (Tilot et al., 2020) and implemented in the LDSC software package (Finucane et al., 2015). Code used to perform analyses is available at https://bitbucket.org/jasonlouisstein/enigmaevolma6/src/master/. Genomic regions that underwent rapid change on the human lineage (human accelerated regions, HARs) were combined from several sources (Capra et al., 2013). Other annotations obtained from previous work included BED files listing fetal brain enhancer elements not found in macaques or mice (Reilly et al., 2015), a refined list of SNPs gained through introgression with Neanderthals (Simonti et al., 2016), genomic regions depleted of introgressed Neanderthal DNA (Vernot et al., 2016), and ancient selective sweep regions identified using extended lineage sorting (Peyrégne et al., 2017). A summary of all annotations can be found in Supplementary Table 1. Enrichment of heritability within HARs, selective sweep regions, Neanderthal-introgressed SNPs, and Neanderthal-depleted regions was controlled for the baselineLD v2 model from the original LDSC study (Finucane et al., 2015). Heritability enrichment in fetal brain HGEs was controlled for both the baseline model and a set of fetal brain active regulatory elements (E081) from the Epigenomics Roadmap resource. Active regulatory elements were defined using chromHMM (Ernst & Kellis, 2012) marks from the 15 state models including all the following annotations: 1_TssA, 2_TssAFlnk, and 7_Enh, 6_EnhG. Enrichment was defined as the proportion of SNP heritability contained in that category divided by the proportion of SNPs in that category. The p-values for heritability estimate and heritability enrichment were Bonferroni corrected for the twenty-two brain sulci tested.

## Results

We examined the contribution of each set of evolutionary annotations to the SNP-based heritability of the sulcal opening and depth of twenty-two brain sulci. The total SNP-based heritability of sulcal opening was significant for all twenty-two brain sulci ranging from 0.08 to 0.21 (Table 1, Figure 1). We found a significant positive enrichment of heritability in sulcal opening in the left (Enrichment = 16.54, Bonferroni corrected *P*-value = 0.040) and right calloso-marginal posterior fissures (Enrichment = 22.44, Bonferroni corrected *P*-value = 0.028) and in the right central sulcus (Enrichment = 19.33, Bonferroni corrected *P*-value = 0.019) with the Human Gained Enhancers (HGEs) active at 7 weeks postconception (Table 1, Figure 2). We did not find significant positive enrichment of heritability for the sulcal opening or depth of the twenty-two brain sulci for any other evolution annotations that we tested: Human Gained Enhancers (active at 8.5 and 12 weeks postconception), human accelerated regions, selective sweeps and Neanderthal introgressed or depleted regions (See Supplementary Tables 2 to 17).

**Table 1.** SNP-based heritability and enrichment in heritability of sulcal opening for Human Gained Enhancers - 7pcw

| Sulcus | Hemisphere | h<sup>2</sup> |  |  |  | Enrichment |  |  |
| --- | --- | --- | --- | --- | --- | --- | --- | --- |
|  |  | Estimate | SE | X<sup>2</sup> | P-value* | Estimate | SE | P-value* |
| FCLp | left | 0.21 | 0.027 | 58.39 | 4.7e-13 | 4.38 | 3.05 | 1 |
| FCLp | right | 0.17 | 0.027 | 41.02 | 3.3e-09 | 10.04 | 4.32 | 0.93 |
| FCMant | left | 0.08 | 0.024 | 11.75 | 0.013 | 8.39 | 10.03 | 1 |
| FCMant | right | 0.09 | 0.025 | 12.98 | 0.0069 | 2.27 | 8.92 | 1 |
| FCMpost | left | 0.15 | 0.025 | 34.85 | 7.8e-08 | 16.54 | 5.03 | 0.04 |
| FCMpost | right | 0.12 | 0.024 | 24.49 | 1.6e-05 | 22.44 | 6.87 | 0.028 |
| FIP | left | 0.16 | 0.028 | 32.43 | 2.7e-07 | 10.16 | 5.16 | 1 |
| FIP | right | 0.15 | 0.026 | 32.73 | 2.3e-07 | 8.58 | 5.27 | 1 |
| SC | left | 0.17 | 0.027 | 38.37 | 1.3e-08 | 17.98 | 5.55 | 0.073 |
| SC | right | 0.16 | 0.029 | 32.18 | 3.1e-07 | 19.33 | 5.50 | 0.019 |
| SFinf | left | 0.16 | 0.024 | 48.46 | 7.4e-11 | 9.14 | 4.22 | 1 |
| SFinf | right | 0.17 | 0.025 | 48.44 | 7.5e-11 | 11.26 | 4.70 | 0.67 |
| SFsup | left | 0.17 | 0.025 | 47.57 | 1.2e-10 | 11.72 | 4.09 | 0.24 |
| SFsup | right | 0.18 | 0.028 | 39.85 | 6e-09 | 12.50 | 4.16 | 0.11 |
| STiant | left | 0.10 | 0.024 | 18.55 | 0.00036 | 7.13 | 5.30 | 1 |
| STiant | right | 0.12 | 0.024 | 25.74 | 8.6e-06 | 16.13 | 9.54 | 1 |
| STipost | left | 0.13 | 0.023 | 29.74 | 1.1e-06 | -2.37 | 8.03 | 1 |
| STipost | right | 0.08 | 0.021 | 15.46 | 0.0019 | 2.68 | 6.34 | 1 |
| STs | left | 0.09 | 0.025 | 14.24 | 0.0035 | 8.40 | 5.10 | 1 |
| STs | right | 0.12 | 0.026 | 20.45 | 0.00013 | 11.97 | 5.23 | 0.73 |
| SsP | left | 0.16 | 0.026 | 37.23 | 2.3e-08 | 10.22 | 7.21 | 1 |
| SsP | right | 0.14 | 0.027 | 26.96 | 4.6e-06 | -0.64 | 5.86 | 1 |

"\*": p-value Bonferroni corrected for the number of sulci. h<sup>2</sup>: SNP-based heritability, X<sup>2</sup>: Chi-squared, SE: Standard Error. FCLp: posterior lateral fissure; FCMant: calloso-marginal anterior fissure; FCMpost: calloso-marginal posterior fissure; FIP: intraparietal sulcus; SC: central sulcus; SFinf: inferior frontal sulcus; SFsup: superior frontal sulcus; STiant: anterior inferior temporal sulcus; STipost: posterior inferior temporal sulcus; STs: superior temporal sulcus; SsP: sub-parietal sulcus.

**Figure 2.**
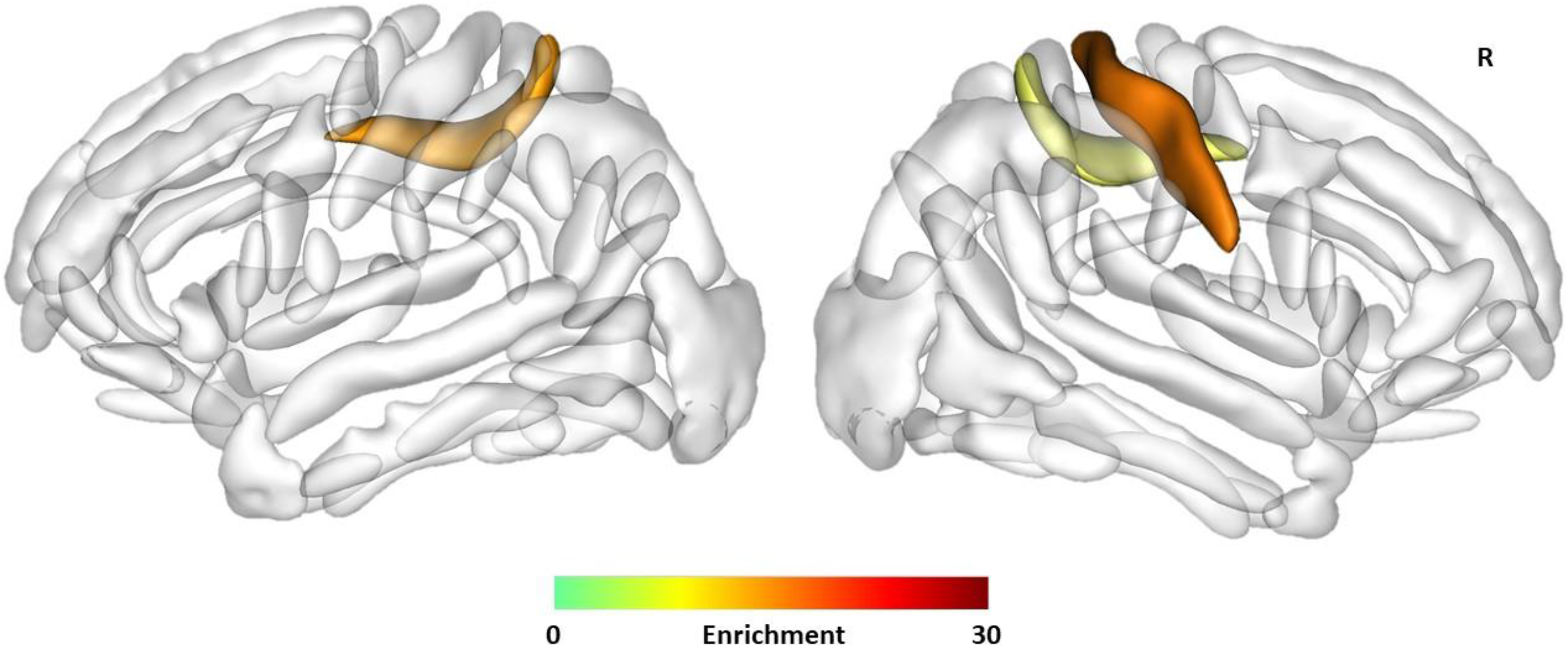
Enrichment of heritability of sulcal opening in the left (Enrichment = 16.54, Bonferroni corrected *P*-value = 0.040) and right calloso-marginal posterior fissures (Enrichment = 22.44, Bonferroni corrected *P*-value = 0.028) and in the right central sulcus (Enrichment = 19.33, Bonferroni corrected *P*-value = 0.019) with the Human Gains Enhancers (HGEs) active at 7 weeks postconception. The sulci are displayed using the statistical probability anatomy map (SPAM) representation, which represents the average sulci shape and position on the reference base of the Brainvisa sulci extraction pipeline (Perrot et al. 2011), R: Right

## Discussion

Using different genomic annotations relevant to human evolution over the last 30 million years, we found that genetic variation within human gained enhancers that are active during fetal development explained more trait heritability than expected for the left and right calloso-marginal posterior fissures and the right central sulcus in adults.

During primate evolution, the formation of gyri and sulci via cortical folding has shown increases of complexity in a structured manner. Cortical folding under the influence of genetic and activity-driven processes (Borrell, 2018) could result from different mechanical forces such as axonal tension, radial expansion or differential tangential expansion (Ronan & Fletcher, 2015). It has also been proposed that such mechanical stresses could in turn affect gene expression and cell behavior during brain development (Foubet et al., 2019). This complex gyrification of the human brain has provided more cortical surface and more space for grey matter cell bodies, neuropil and glial cells (Namba & Huttner, 2017), which might have contributed to cognitive changes in modern humans related to distinct traits like creativity and language.

From the anatomical point of view, the central sulcus is one of the major sulci in the brain that has been of particular interest in the evolution in primates. Located in the middle of sensory-motor systems, it has been related to the fine tuning of the motor system in apes and specifically to the increasing motor control of the hand. Relative to the total cortical surface, the central sulcus surface has increased along the phylogenetic timeline of primates/apes, based on comparative analyses of Old World monkeys (common ancestor ≈ 30 Mya), lesser apes (≈ 20-15 Mya) and great apes (≈ 15-10 Mya), reaching its maximum in the orangutan and gorilla and then decreasing in chimpanzee and human (Hopkins et al., 2014). Moreover, great apes and humans show a distinct dorsal ventral pattern in the central sulcus presumably due to the folding over the buried “pli de passage fronto parietal moyen” (PPFM) referred to as the motor-hand area or hand knob (J.-F. Mangin et al., 2019). In humans, the PPFM gyrus has been hypothesized to relate to adaptations for bipedalism and increased use of the hands for tool-use and other manual functions (Hopkins et al., 2014).

The calloso-marginal posterior fissure is the posterior part of the marginal sulcus and borders the paracentral lobule on the medial surface of the cortex where the primary sensory and motor cortices merge. This region has been linked to the control of the hind limbs and lower extremities (i.e. leg and foot) (Liao et al., 2016). It has been suggested that anatomical changes within this region could be relevant to the evolution of locomotion in primates including different types of quadrupedalism, leaping, brachiation and, in rare cases, bipedalism (Dial et al., 2015). For instance, such changes could be related to the transition from arboreal to terrestrial quadrupedalism mostly found in Old World monkeys during the Oligocene period around 30 million years ago (Napier, 1967).

From the genetic point of view, human gained enhancers are regulatory genetic elements that emerged since our last common ancestor with Old World monkeys about 30 million years ago (Reilly et al., 2015). These elements were detected by comparing regulatory elements between mice, macaques, and humans during corticogenesis. Specifically, researchers profiled post-translational histone modifications (H3K27ac and H3K4me2) to map active promoters and enhancers in mice, macaques and humans, and to identify increases in gene activity in human cortex from 7 to 12 post conception weeks (Reilly et al., 2015). This work revealed human gained activity in gene modules important for neuronal proliferation, migration, and cortical-map organization. In the present study, our results suggest that common genetic variations in living human populations within these regulatory elements are related to sulcal morphology. By doing so, it gives us some insight into possible biological functions gained during human evolution. Looking at non-brain features, human gained enhancers have been associated with human limb development (Cotney et al., 2013), and are argued to be important for specific changes in hindlimb morphology such as the characteristic inflexibility and shortened digits of the human foot (Prabhakar et al., 2008). Given that human gained enhancers are active in the brain and that their genetic variations are related to sulcal features in the dorsal sensory-motor system, we may speculate that they played a role in the evolution of locomotion in primates and contributed to adaptations to our unique form bipedalism.

In this study, we focused on sulcal opening and depth features to assess sulcal morphology instead of cortical measures such as thickness and surface area. Sulcal morphometry may be more related to differential neuronal proliferation (Reillo et al., 2011) and to axonal fiber constraints (Van Essen, 1997). On the other hand, cortical thickness seems more related to cellular and laminar organization of the cortical mantle (Wagstyl et al., 2015), while surface area is driven by the number of radial columns perpendicular to the pial surface (Rakic, 2009). This distinction may explain why our analyses of human gained enhancers and sulcal phenotypes highlighted different cortical regions from those for a similar analysis focused on surface area (Tilot et al., 2020), as these different biological mechanisms are not controlled by the same genetic factors. One particular result is that enrichment of heritability is only present for sulcal opening and not for sulcal depth. Sulcal opening being more able to capture degenerative processes (Kochunov et al., 2005), it is interesting to see this trait is also more able to capture genetic variation. In a recent study of more than 9,000 subjects, sulcal opening was found to be the most heritable and reliable sulcal metric (Pizzagalli et al., 2020), which could explain the strong relationship between this metric and the genomic annotations.

Our findings should be considered in light of some limitations. We limited our study to genetic annotations over a selection of broadly defined periods of primate evolutionary history. Other approaches could be used with better temporal resolution and more comprehensive temporal distribution along lineages that led to emergence of Homo sapiens (Albers & McVean, 2020; Andirkó et al., 2021). Moreover, with our methodology we assessed only genetic variations that are polymorphic in current human populations. This study design does not allow assessment of effects of ancestral changes that are now fixed (non-variant) in modern humans, although such variations are also very likely to have contributed to the evolution of cortical gyrification. Finally, we did not interpret significant negative enrichments of heritability for the Neanderthal introgressed SNPs (See Table S8). Indeed, the LDSC software used to estimate partitioned heritability allows for negative enrichment, but these values are unexpected as there are no biologically plausible mechanisms by which such directions of effect could occur. The significance of these negative estimates could be due to the rather rare genetic variations in modern humans for this specific annotation. In this case, the effect sizes within this annotation could be not normally distributed producing model instability and making enrichment values uninterpretable.

This study on cortical gyrification expands beyond prior work showing genetic variations within human gained enhancers have effects on surface area of multiple cortical regions such as the inferior frontal gyrus, a region known for its involvement in language processing (Tilot et al., 2020). In our study, we showed that genetic variations in genomic regions that have shown different regulatory activity over the last 30 million years of human evolution may have also impacted on the shape of different human sulci, albeit in different regions of the cortex. Based on the literature, the relationship between sulci, namely the central sulcus and calloso-marginal posterior fissure, and the human gained enhancers may possibly relate to the evolution of locomotion in primates and bipedalism in human, generating hypotheses for further investigation. Future studies may help elaborate further on the neurobiological substrates of such evolution patterns.

## Acknowledgement

This research has been conducted using data from UK Biobank, a major biomedical database” and where appropriate, include a link to the UK Biobank website: www.ukbiobank.ac.uk

