## Supplementary Material for "Genetic variations within human gained enhancer elements affect human brain sulcal morphology"

**Table S1: Characteristics of human-evolution focused genome annotations**

| Name | # Regions | Total Size (kb) | # SNPs | Average MAF | Reference |
|---|---|---|---|---|---|
| **Human Accelerated Regions** | 2,737 | 702 | 1.874 | 0.213 | Capra, Philos Trans R Soc Lond B Biol Sci., 2013 |
| **Human Gained Enhancers - 7pcw** | 7,742 | 22,195 | 57,059 | 0.205 | Reilly, Science, 2015 |
| **Human Gained Enhancers - 8.5pcw** | 5,101 | 12,277 | 29.985 | 0.203 | Reilly, Science, 2015 |
| **Human Gained Enhancers - 12pcw (Frontal)** | 3,110 | 8,079 | 20,741 | 0.203 | Reilly, Science, 2015 |
| **Human Gained Enhancers - 12pcw (Occipital)** | 4,994 | 12,297 | 29,344 | 0.203 | Reilly, Science, 2015 |
| **Neandertal Introgression SNPs** | 5,851 | 722,954 | 24,331 | 0.115 | Simonti, Science, 2016 |
| **Neandertal Lineage Depleted Regions** | 6 | 84,440 | 175,377 | 0.208 | Vernot, Science, 2016 |
| **Selective Sweeps** | 314 | 19,064 | 23,744 | 0.196 | Peyrégne, Genome Res, 2017 |

Kb: Kilobase, MAF: Minor Allele Frequency, pcw: post conception weeks, SNP: single nucleotide polymorphism.

**Table S2. Enrichment in heritability of sulcal opening for Human Gained Enhancers - 8.5pcw**

| Sulcus | Hemisphere | Enrichment | Enrichment SE | P-value* |
|---|---|---|---|---|
| FCLp | left | 7.34 | 5.95 | 1 |
| FCLp | right | 15.42 | 7.61 | 1 |
| FCMant | left | -1.20 | 16.53 | 1 |
| FCMant | right | -9.75 | 16.11 | 1 |
| FCMpost | left | 26.01 | 10.05 | 0.32 |
| FCMpost | right | 21.62 | 11.99 | 1 |
| FIP | left | 13.50 | 8.49 | 1 |
| FIP | right | 5.64 | 8.83 | 1 |
| SC | left | 14.90 | 9.66 | 1 |
| SC | right | 14.50 | 8.93 | 1 |
| SFinf | left | 17.46 | 7.07 | 0.53 |
| SFinf | right | 21.58 | 7.68 | 0.14 |
| SFsup | left | 6.83 | 7.63 | 1 |
| SFsup | right | 10.33 | 6.75 | 1 |
| STiant | left | 17.34 | 9.34 | 1 |
| STiant | right | 29.00 | 16.77 | 1 |
| STipost | left | 3.65 | 14.15 | 1 |
| STipost | right | 10.84 | 10.36 | 1 |
| STs | left | 13.49 | 8.14 | 1 |
| STs | right | 28.37 | 9.68 | 0.077 |
| SsP | left | 7.63 | 11.32 | 1 |
| SsP | right | -6.40 | 9.16 | 1 |

"*": p-value Bonferroni corrected for the number of sulci. SE: Standard Error.

**Table S3. Enrichment in heritability of sulcal opening for Human Gained Enhancers - 12pcw (Frontal)**

| Sulcus | Hemisphere | Enrichment | Enrichment SE | P-value* |
|---|---|---|---|---|
| FCLp | left | 6.52 | 7.30 | 1 |
| FCLp | right | 13.69 | 9.05 | 1 |
| FCMant | left | -8.67 | 17.73 | 1 |
| FCMant | right | -7.81 | 19.34 | 1 |
| FCMpost | left | 32.37 | 13.58 | 0.58 |
| FCMpost | right | 18.82 | 14.93 | 1 |
| FIP | left | 12.66 | 12.21 | 1 |
| FIP | right | -2.77 | 11.38 | 1 |
| SC | left | 9.04 | 10.36 | 1 |
| SC | right | 8.23 | 10.54 | 1 |
| SFinf | left | 16.24 | 9.37 | 1 |
| SFinf | right | 29.17 | 9.55 | 0.058 |
| SFsup | left | 6.03 | 9.69 | 1 |
| SFsup | right | 4.95 | 9.32 | 1 |
| STiant | left | 8.11 | 10.62 | 1 |
| STiant | right | 29.58 | 19.78 | 1 |
| STipost | left | 3.28 | 16.80 | 1 |
| STipost | right | -4.12 | 14.93 | 1 |
| STs | left | 11.60 | 9.95 | 1 |
| STs | right | 20.64 | 11.66 | 1 |
| SsP | left | 14.39 | 14.62 | 1 |
| SsP | right | -5.55 | 14.67 | 1 |

"*": p-value Bonferroni corrected for the number of sulci. SE: Standard Error.

**Table S4. Enrichment in heritability of sulcal opening for Human Gained Enhancers - 12pcw (Occipital)**

| Sulcus | Hemisphere | Enrichment | Enrichment SE | P-value* |
|---|---|---|---|---|
| FCLp | left | 4.32 | 6.37 | 1 |
| FCLp | right | 12.21 | 7.51 | 1 |
| FCMant | left | -9.74 | 15.99 | 1 |
| FCMant | right | 7.32 | 14.74 | 1 |
| FCMpost | left | 25.63 | 11.35 | 0.75 |
| FCMpost | right | 18.39 | 11.47 | 1 |
| FIP | left | 14.05 | 8.46 | 1 |
| FIP | right | 4.65 | 8.75 | 1 |
| SC | left | 13.54 | 7.63 | 1 |
| SC | right | 10.50 | 8.49 | 1 |
| SFinf | left | 18.98 | 8.85 | 0.99 |
| SFinf | right | 14.30 | 7.73 | 1 |
| SFsup | left | 21.50 | 10.56 | 1 |
| SFsup | right | 13.16 | 7.80 | 1 |
| STiant | left | 8.58 | 11.76 | 1 |
| STiant | right | 32.14 | 20.52 | 1 |
| STipost | left | 14.72 | 15.68 | 1 |
| STipost | right | 9.27 | 12.92 | 1 |
| STs | left | 12.94 | 8.65 | 1 |
| STs | right | 18.89 | 9.73 | 1 |
| SsP | left | 18.23 | 13.06 | 1 |
| SsP | right | -7.98 | 10.98 | 1 |

"*": p-value Bonferroni corrected for the number of sulci. SE: Standard Error.

**Table S5. Enrichment in heritability of sulcal opening for Selective Sweeps**

| Sulcus | Hemisphere | Enrichment | Enrichment SE | P-value* |
|---|---|---|---|---|
| FCLp | left | 1.74 | 2.62 | 1 |
| FCLp | right | 2.48 | 3.73 | 1 |
| FCMant | left | 9.02 | 7.96 | 1 |
| FCMant | right | 12.28 | 11.27 | 1 |
| FCMpost | left | 9.28 | 7.46 | 1 |
| FCMpost | right | -1.74 | 4.97 | 1 |
| FIP | left | 11.58 | 7.88 | 1 |
| FIP | right | 10.03 | 6.77 | 1 |
| SC | left | 5.77 | 5.72 | 1 |
| SC | right | 1.29 | 3.46 | 1 |
| SFinf | left | -3.48 | 3.51 | 1 |
| SFinf | right | 2.95 | 3.07 | 1 |
| SFsup | left | 3.50 | 5.16 | 1 |
| SFsup | right | 5.67 | 5.15 | 1 |
| STiant | left | 0.51 | 3.51 | 1 |
| STiant | right | 7.30 | 6.77 | 1 |
| STipost | left | 11.61 | 8.28 | 1 |
| STipost | right | 12.67 | 6.75 | 1 |
| STs | left | 1.98 | 3.31 | 1 |
| STs | right | 2.96 | 3.80 | 1 |
| SsP | left | 10.41 | 5.99 | 1 |
| SsP | right | 5.62 | 3.49 | 1 |

"*": p-value Bonferroni corrected for the number of sulci. SE: Standard Error.

**Table S6. Enrichment in heritability of sulcal opening for Human Accelerated Regions**

| Sulcus | Hemisphere | Enrichment | Enrichment SE | P-value* |
|---|---|---|---|---|
| FCLp | left | 1.64 | 66.73 | 1 |
| FCLp | right | 42.55 | 84.87 | 1 |
| FCMant | left | -11.76 | 130.90 | 1 |
| FCMant | right | -121.02 | 129.23 | 1 |
| FCMpost | left | 43.29 | 108.66 | 1 |
| FCMpost | right | -132.44 | 122.24 | 1 |
| FIP | left | 13.07 | 95.42 | 1 |
| FIP | right | -65.46 | 91.38 | 1 |
| SC | left | -72.29 | 81.07 | 1 |
| SC | right | -71.61 | 78.83 | 1 |
| SFinf | left | -83.89 | 83.52 | 1 |
| SFinf | right | 70.90 | 70.24 | 1 |
| SFsup | left | -116.62 | 69.09 | 1 |
| SFsup | right | -112.90 | 63.97 | 1 |
| STiant | left | 54.75 | 90.20 | 1 |
| STiant | right | 102.09 | 152.88 | 1 |
| STipost | left | 90.81 | 164.24 | 1 |
| STipost | right | -4.66 | 116.59 | 1 |
| STs | left | 60.45 | 98.57 | 1 |
| STs | right | -13.30 | 91.25 | 1 |
| SsP | left | -132.21 | 146.24 | 1 |
| SsP | right | -82.38 | 89.05 | 1 |

"*": p-value Bonferroni corrected for the number of sulci. SE: Standard Error.

**Table S7. Enrichment in heritability of sulcal opening for Neandertal Lineage Depleted Regions**

| Sulcus | Hemisphere | Enrichment | Enrichment SE | P-value* |
|---|---|---|---|---|
| FCLp | left | 1.37 | 0.79 | 1 |
| FCLp | right | 1.22 | 0.79 | 1 |
| FCMant | left | 1.09 | 1.82 | 1 |
| FCMant | right | 2.05 | 1.24 | 1 |
| FCMpost | left | 0.25 | 0.85 | 1 |
| FCMpost | right | 1.04 | 0.94 | 1 |
| FIP | left | 0.35 | 1.06 | 1 |
| FIP | right | 0.70 | 1.02 | 1 |
| SC | left | 0.15 | 0.53 | 1 |
| SC | right | 0.35 | 0.58 | 1 |
| SFinf | left | 0.79 | 0.65 | 1 |
| SFinf | right | 0.39 | 0.56 | 1 |
| SFsup | left | 0.39 | 0.59 | 1 |
| SFsup | right | 0.80 | 0.73 | 1 |
| STiant | left | 0.70 | 0.57 | 1 |
| STiant | right | 2.20 | 1.13 | 1 |
| STipost | left | -0.81 | 0.81 | 0.49 |
| STipost | right | -0.11 | 0.67 | 1 |
| STs | left | 1.11 | 0.60 | 1 |
| STs | right | 0.96 | 0.64 | 1 |
| SsP | left | 0.15 | 0.96 | 1 |
| SsP | right | 0.49 | 1.00 | 1 |

"*": p-value Bonferroni corrected for the number of sulci. SE: Standard Error.

**Table S8. Enrichment in heritability of sulcal opening for Neandertal Introgression SNPs**

| Sulcus | Hemisphere | Enrichment | Enrichment SE | P-value* |
|---|---|---|---|---|
| FCLp | left | -8.21 | 2.18 | 0.00021 |
| FCLp | right | -7.16 | 2.46 | 0.017 |
| FCMant | left | 3.13 | 5.58 | 1 |
| FCMant | right | -5.71 | 4.41 | 1 |
| FCMpost | left | -7.58 | 2.85 | 0.098 |
| FCMpost | right | -12.22 | 3.96 | 0.0022 |
| FIP | left | -4.18 | 2.83 | 1 |
| FIP | right | -4.82 | 2.60 | 0.74 |
| SC | left | -6.79 | 2.53 | 0.04 |
| SC | right | -7.07 | 2.54 | 0.033 |
| SFinf | left | -3.13 | 2.94 | 1 |
| SFinf | right | -2.66 | 2.30 | 1 |
| SFsup | left | -6.11 | 2.55 | 0.066 |
| SFsup | right | -2.83 | 2.62 | 1 |
| STiant | left | 1.15 | 3.63 | 1 |
| STiant | right | 0.47 | 5.91 | 1 |
| STipost | left | -10.67 | 4.84 | 0.14 |
| STipost | right | -2.73 | 4.29 | 1 |
| STs | left | -7.69 | 2.77 | 0.017 |
| STs | right | -2.80 | 3.73 | 1 |
| SsP | left | -2.07 | 4.63 | 1 |
| SsP | right | 2.18 | 3.62 | 1 |

"*": p-value Bonferroni corrected for the number of sulci. SE: Standard Error.

**Table S9. SNP-based heritability estimates of sulcal depth**

| Sulcus | Hemisphere | $h^2$ | $h^2$ SE | $X^2$ | P-value* |
|---|---|---|---|---|---|
| FCLp | left | 0.03 | 0.027 | 1.41 | 1 |
| FCLp | right | 0.07 | 0.021 | 10.38 | 0.028 |
| FCMant | left | 0.03 | 0.022 | 1.88 | 1 |
| FCMant | right | 0.05 | 0.022 | 4.99 | 0.56 |
| FCMpost | left | 0.13 | 0.029 | 19.28 | 0.00025 |
| FCMpost | right | 0.10 | 0.025 | 16.46 | 0.0011 |
| FIP | left | 0.09 | 0.026 | 12.92 | 0.0072 |
| FIP | right | 0.13 | 0.024 | 31.56 | 4.3e-07 |
| SC | left | 0.09 | 0.022 | 18.48 | 0.00038 |
| SC | right | 0.10 | 0.023 | 18.33 | 0.00041 |
| SFinf | left | 0.10 | 0.022 | 21.37 | 8.3e-05 |
| SFinf | right | 0.08 | 0.025 | 9.38 | 0.048 |
| SFsup | left | 0.06 | 0.027 | 5.34 | 0.46 |
| SFsup | right | 0.05 | 0.024 | 3.92 | 1 |
| SsP | left | 0.08 | 0.026 | 9.38 | 0.048 |
| SsP | right | 0.03 | 0.023 | 1.36 | 1 |
| STiant | left | 0.10 | 0.022 | 20.05 | 0.00017 |
| STiant | right | 0.10 | 0.025 | 15.95 | 0.0014 |
| STipost | left | 0.04 | 0.022 | 2.74 | 1 |
| STipost | right | 0.08 | 0.024 | 10.93 | 0.021 |
| STs | left | 0.10 | 0.024 | 16.92 | 0.00086 |
| STs | right | 0.08 | 0.025 | 9.14 | 0.055 |

"*": p-value Bonferroni corrected for the number of sulci. $h^2$: SNP-based heritability, $X^2$: Chi-squared, SE: Standard Error.

**Table S10. Enrichment in heritability of sulcal depth for Human Gained Enhancers - 7pcw**

| Sulcus | Hemisphere | Enrichment | Enrichment SE | P-value* |
|---|---|---|---|---|
| FCLp | left | -18.77 | 29.60 | 1 |
| FCLp | right | -6.07 | 10.74 | 1 |
| FCMant | left | -53.27 | 53.83 | 0.55 |
| FCMant | right | -13.77 | 14.14 | 1 |
| FCMpost | left | -7.20 | 6.35 | 1 |
| FCMpost | right | 14.04 | 8.90 | 1 |
| FIP | left | 10.33 | 7.61 | 1 |
| FIP | right | 5.32 | 4.72 | 1 |
| SC | left | 16.74 | 8.21 | 1 |
| SC | right | -1.19 | 6.34 | 1 |
| SFinf | left | 14.33 | 5.84 | 0.52 |
| SFinf | right | 13.64 | 12.43 | 1 |
| SFsup | left | 25.75 | 13.26 | 0.65 |
| SFsup | right | 27.05 | 15.78 | 1 |
| STiant | left | -2.45 | 7.89 | 1 |
| STiant | right | 14.18 | 7.21 | 1 |
| STipost | left | 3.94 | 19.57 | 1 |
| STipost | right | -6.46 | 9.58 | 1 |
| STs | left | 9.14 | 6.91 | 1 |
| STs | right | -2.44 | 9.61 | 1 |
| SsP | left | 7.81 | 7.89 | 1 |
| SsP | right | 39.47 | 28.33 | 1 |

"*": p-value Bonferroni corrected for the number of sulci. SE: Standard Error.

**Table S11. Enrichment in heritability of sulcal depth for Human Gained Enhancers - 8.5pcw**

| Sulcus | Hemisphere | Enrichment | Enrichment SE | P-value* |
|---|---|---|---|---|
| FCLp | left | -39.42 | 48.23 | 1 |
| FCLp | right | 0.83 | 17.97 | 1 |
| FCMant | left | -71.39 | 60.92 | 0.78 |
| FCMant | right | -2.57 | 21.77 | 1 |
| FCMpost | left | 7.16 | 9.83 | 1 |
| FCMpost | right | -5.05 | 13.34 | 1 |
| FIP | left | 5.37 | 13.37 | 1 |
| FIP | right | 13.12 | 11.14 | 1 |
| SC | left | 6.35 | 14.73 | 1 |
| SC | right | -21.97 | 12.84 | 1 |
| SFinf | left | 14.82 | 13.68 | 1 |
| SFinf | right | -4.64 | 19.47 | 1 |
| SFsup | left | 35.88 | 20.78 | 1 |
| SFsup | right | 14.44 | 28.22 | 1 |
| STiant | left | -3.17 | 12.50 | 1 |
| STiant | right | 26.53 | 15.33 | 1 |
| STipost | left | -28.50 | 44.68 | 1 |
| STipost | right | -8.75 | 15.75 | 1 |
| STs | left | 16.63 | 10.75 | 1 |
| STs | right | -1.95 | 17.86 | 1 |
| SsP | left | 12.75 | 15.19 | 1 |
| SsP | right | 46.78 | 45.44 | 1 |

"*": p-value Bonferroni corrected for the number of sulci. SE: Standard Error.

**Table S12. Enrichment in heritability of sulcal depth for Human Gained Enhancers - 12pcw (Frontal)**

| Sulcus | Hemisphere | Enrichment | Enrichment SE | P-value* |
|---|---|---|---|---|
| FCLp | left | -93.26 | 87.80 | 0.45 |
| FCLp | right | -17.20 | 22.54 | 1 |
| FCMant | left | -39.96 | 47.62 | 1 |
| FCMant | right | 5.27 | 26.37 | 1 |
| FCMpost | left | 2.72 | 13.65 | 1 |
| FCMpost | right | -15.06 | 13.52 | 1 |
| FIP | left | -19.07 | 17.62 | 1 |
| FIP | right | -2.59 | 12.92 | 1 |
| SC | left | 22.31 | 15.16 | 1 |
| SC | right | -25.28 | 14.70 | 1 |
| SFinf | left | -16.78 | 16.09 | 1 |
| SFinf | right | 20.07 | 21.32 | 1 |
| SFsup | left | 61.55 | 34.81 | 0.22 |
| SFsup | right | 41.72 | 30.23 | 1 |
| STiant | left | 12.34 | 14.38 | 1 |
| STiant | right | 33.77 | 19.54 | 1 |
| STipost | left | -32.08 | 52.53 | 1 |
| STipost | right | -1.50 | 18.27 | 1 |
| STs | left | 10.31 | 16.26 | 1 |
| STs | right | -39.49 | 25.03 | 0.63 |
| SsP | left | 21.93 | 18.82 | 1 |
| SsP | right | 35.53 | 71.25 | 1 |

"*": p-value Bonferroni corrected for the number of sulci. SE: Standard Error.

**Table S13. Enrichment in heritability of sulcal depth for Human Gained Enhancers - 12pcw (Occipital)**

| Sulcus | Hemisphere | Enrichment | Enrichment SE | P-value* |
|---|---|---|---|---|
| FCLp | left | -57.19 | 64.14 | 1 |
| FCLp | right | -9.50 | 19.30 | 1 |
| FCMant | left | 4.97 | 32.77 | 1 |
| FCMant | right | 8.83 | 21.57 | 1 |
| FCMpost | left | 2.21 | 10.87 | 1 |
| FCMpost | right | -4.16 | 13.36 | 1 |
| FIP | left | 8.35 | 14.05 | 1 |
| FIP | right | 4.40 | 10.54 | 1 |
| SC | left | 1.22 | 14.91 | 1 |
| SC | right | -9.89 | 14.59 | 1 |
| SFinf | left | 8.17 | 14.91 | 1 |
| SFinf | right | 4.95 | 21.52 | 1 |
| SFsup | left | 30.52 | 22.91 | 1 |
| SFsup | right | 46.62 | 29.64 | 1 |
| STiant | left | 14.09 | 13.58 | 1 |
| STiant | right | 43.88 | 16.07 | 0.12 |
| STipost | left | -95.73 | 80.43 | 0.32 |
| STipost | right | -14.26 | 15.37 | 1 |
| STs | left | -7.87 | 15.83 | 1 |
| STs | right | -47.87 | 25.52 | 0.079 |
| SsP | left | 17.08 | 16.60 | 1 |
| SsP | right | 24.72 | 49.60 | 1 |

"*": p-value Bonferroni corrected for the number of sulci. SE: Standard Error.

**Table S14. Enrichment in heritability of sulcal depth for Selective Sweeps**

| Sulcus | Hemisphere | Enrichment | Enrichment SE | P-value* |
|---|---|---|---|---|
| FCLp | left | -14.28 | 23.22 | 1 |
| FCLp | right | 4.91 | 6.44 | 1 |
| FCMant | left | 33.39 | 50.11 | 1 |
| FCMant | right | -3.04 | 13.97 | 1 |
| FCMpost | left | 7.93 | 5.60 | 1 |
| FCMpost | right | 3.73 | 5.14 | 1 |
| FIP | left | 1.01 | 4.93 | 1 |
| FIP | right | 8.93 | 4.96 | 1 |
| SC | left | 1.76 | 5.24 | 1 |
| SC | right | 9.05 | 6.15 | 1 |
| SFinf | left | -0.23 | 4.01 | 1 |
| SFinf | right | -5.12 | 5.27 | 1 |
| SFsup | left | -2.56 | 8.46 | 1 |
| SFsup | right | 5.99 | 10.68 | 1 |
| STiant | left | 9.96 | 7.80 | 1 |
| STiant | right | 5.79 | 4.90 | 1 |
| STipost | left | 0.05 | 9.58 | 1 |
| STipost | right | -1.32 | 4.78 | 1 |
| STs | left | -2.33 | 2.96 | 1 |
| STs | right | 3.70 | 5.40 | 1 |
| SsP | left | 3.38 | 6.23 | 1 |
| SsP | right | -26.68 | 54.06 | 1 |

"*": p-value Bonferroni corrected for the number of sulci. SE: Standard Error.

**Table S15. Enrichment in heritability of sulcal depth for Human Accelerated Regions**

| Sulcus | Hemisphere | Enrichment | Enrichment SE | P-value* |
|---|---|---|---|---|
| FCLp | left | 105.31 | 435.74 | 1 |
| FCLp | right | -276.77 | 167.69 | 1 |
| FCMant | left | -456.20 | 785.94 | 1 |
| FCMant | right | -304.77 | 408.01 | 1 |
| FCMpost | left | 47.22 | 97.83 | 1 |
| FCMpost | right | 52.58 | 129.47 | 1 |
| FIP | left | 67.88 | 155.02 | 1 |
| FIP | right | -72.97 | 79.08 | 1 |
| SC | left | -54.59 | 129.25 | 1 |
| SC | right | 164.17 | 129.29 | 1 |
| SFinf | left | 66.86 | 114.03 | 1 |
| SFinf | right | 144.35 | 174.37 | 1 |
| SFsup | left | 46.96 | 268.13 | 1 |
| SFsup | right | -281.88 | 356.12 | 1 |
| STiant | left | 56.91 | 124.05 | 1 |
| STiant | right | 17.42 | 116.03 | 1 |
| STipost | left | 68.59 | 258.35 | 1 |
| STipost | right | 312.66 | 173.67 | 0.87 |
| STs | left | -6.65 | 122.80 | 1 |
| STs | right | -43.01 | 163.89 | 1 |
| SsP | left | -79.45 | 151.57 | 1 |
| SsP | right | 663.75 | 1232.23 | 1 |

"*": p-value Bonferroni corrected for the number of sulci. SE: Standard Error.

**Table S16. Enrichment in heritability of sulcal depth for Neandertal Lineage Depleted Regions**

| Sulcus | Hemisphere | Enrichment | Enrichment SE | P-value* |
|---|---|---|---|---|
| FCLp | left | 7.46 | 9.11 | 0.31 |
| FCLp | right | 0.86 | 1.15 | 1 |
| FCMant | left | 5.77 | 9.82 | 1 |
| FCMant | right | 1.70 | 1.60 | 1 |
| FCMpost | left | 2.19 | 1.34 | 1 |
| FCMpost | right | -0.80 | 1.39 | 1 |
| FIP | left | 1.09 | 1.58 | 1 |
| FIP | right | 1.61 | 0.94 | 1 |
| SC | left | 2.06 | 1.27 | 1 |
| SC | right | 1.76 | 1.32 | 1 |
| SFinf | left | 3.06 | 1.11 | 0.95 |
| SFinf | right | -1.44 | 1.24 | 0.52 |
| SFsup | left | -1.20 | 1.35 | 0.54 |
| SFsup | right | -1.25 | 2.96 | 1 |
| STiant | left | 1.07 | 0.77 | 1 |
| STiant | right | 1.16 | 1.58 | 1 |
| STipost | left | -0.48 | 2.40 | 1 |
| STipost | right | 2.62 | 1.41 | 1 |
| STs | left | 1.64 | 1.32 | 1 |
| STs | right | 2.26 | 1.65 | 1 |
| SsP | left | 0.45 | 1.47 | 1 |
| SsP | right | 6.82 | 13.39 | 1 |

"*": p-value Bonferroni corrected for the number of sulci. SE: Standard Error.

**Table S17. Enrichment in heritability of sulcal depth for Neandertal Introgression SNPs**

| Sulcus | Hemisphere | Enrichment | Enrichment SE | P-value* |
|---|---|---|---|---|
| FCLp | left | 7.22 | 16.93 | 1 |
| FCLp | right | 12.09 | 8.51 | 1 |
| FCMant | left | 14.02 | 27.16 | 1 |
| FCMant | right | 0.37 | 13.54 | 1 |
| FCMpost | left | -2.04 | 3.86 | 1 |
| FCMpost | right | -4.78 | 4.37 | 1 |
| FIP | left | -15.61 | 5.66 | 0.0057 |
| FIP | right | 3.39 | 3.76 | 1 |
| SC | left | 1.35 | 4.20 | 1 |
| SC | right | -3.91 | 4.46 | 1 |
| SFinf | left | 5.92 | 4.61 | 1 |
| SFinf | right | 3.84 | 6.42 | 1 |
| SFsup | left | -7.03 | 8.58 | 1 |
| SFsup | right | -3.54 | 12.76 | 1 |
| STiant | left | -5.19 | 3.77 | 1 |
| STiant | right | -5.32 | 3.76 | 1 |
| STipost | left | -7.52 | 10.99 | 1 |
| STipost | right | 0.23 | 4.35 | 1 |
| STs | left | -5.74 | 4.86 | 1 |
| STs | right | -12.35 | 5.13 | 0.035 |
| SsP | left | -1.73 | 6.02 | 1 |
| SsP | right | 1.11 | 25.78 | 1 |

"*": p-value Bonferroni corrected for the number of sulci. SE: Standard Error.

**Sulci acronyms:** FCLp: posterior lateral fissure; FCMant: calloso-marginal anterior fissure; FCMpost: calloso-marginal posterior fissure; FIP: intraparietal sulcus; SC: central sulcus; SFinf: inferior frontal sulcus; SFsup: superior frontal sulcus; STiant: anterior inferior temporal sulcus; STipost: posterior inferior temporal sulcus; STs: superior temporal sulcus; SsP: sub-parietal sulcus.
